# Double-strand break repair-associated intragenic deletions and tandem duplications suggest the architecture of the repair replication fork

**DOI:** 10.1101/2023.10.09.561461

**Authors:** Simona Dalin, Sophie Webster, Neal Sugawara, Shu Zhang, Qiuqin Wu, Tracy Cui, Victoria Liang, Rameen Beroukhim, James E. Haber

## Abstract

Double-strand break (DSB) repair is associated with a 1000-fold increase in mutations compared to normal replication of the same sequences. In budding yeast, repair of an HO endonuclease-induced DSB at the *MAT*α locus can be repaired by using a homologous, heterochromatic *HMR::Kl-URA3* donor harboring a transcriptionally silenced *URA3* gene, resulting in a *MAT::URA3* (Ura+) repair product where *URA3* is expressed. Repair-associated *ura3-* mutations can be selected by resistance to 5-fluoroorotic acid (FOA). Using this system, we find that a major class of mutations are -1 deletions, almost always in homonucleotide runs, but there are few +1 insertions. In contrast, +1 and -1 insertions in homonucleotide runs are nearly equal among spontaneous mutations. Approximately 10% of repair-associated mutations are interchromosomal template switches (ICTS), even though the *K. lactis URA3* sequence embedded in *HMR* is only 72% identical with *S. cerevisiae ura3-52* sequences on a different chromosome. ICTS events begin and end in regions of short microhomology, averaging 7 bp. Long microhomologies are favored, but some ICTS junctions are as short as 2 bp. Both repair-associated intragenic deletions (IDs) and tandem duplications (TDs) are recovered, with junctions sharing short stretches of, on average, 6 bp of microhomology. Intragenic deletions are more than 5 times more frequent than TDs. IDs have a mean length of 60 bp, but, surprisingly there are almost no deletions shorter than 25 bp. In contrast, TDs average only 12 bp. The usage of microhomologies among intragenic deletions is not strongly influenced by the degree of adjacent homeology. Together, these data provide a picture of the structure of the repair replication fork. We suggest that IDs and TDs occur within the migrating D-loop in which DNA polymerase δ copies the template, where the 3’ end of a partly copied new DNA strand can dissociate and anneal with a single-stranded region of microhomology that lies either in front or behind the 3’ end, within the open structure of a migrating D-loop. Our data suggest that ∼100 bp ahead of the polymerase is “open,” but that part of the repair replication apparatus remains bound in the 25 bp ahead of the newly copied DNA, preventing annealing. In contrast, the template region behind the polymerase appears to be rapidly reannealed, limiting template switching to a very short region.

## Introduction

Double-strand breaks (DSBs) pose an existential threat to cells, which have evolved a variety of repair mechanisms to restore genome integrity. In mammalian cells, where spontaneous DSBs occur multiple times per DNA replication cycle, genes that encode the homologous recombination (HR) repair machinery are essential. In budding yeast, with its much smaller genome, such breaks occur only about one every 10 cell cycles, but when they arise, their repair also requires the homologous recombination proteins such as Rad51 or Rad52 ^1^. The most accurate of the HR mechanisms is gene conversion, in which both ends of a DSB engage with a homologous template – a sister chromatid, a homolog or an ectopic sequence – to patch up the break by a short patch of new DNA synthesis (synthesis dependent strand annealing, SDSA). A related mechanism is break-induced replication (BIR), in which only one end of the DSB successfully engages the homologous template sequences.

DNA synthesis during DSB repair is strikingly different from normal DNA replication. In SDSA, the predominant mechanism employed in mitotic cells (**Figure 1a**), the two strands of the repaired locus are synthesized sequentially and not simultaneously, as they would during normal replication. Moreover, the newly copied DNA is not semi-conservatively replicated; instead, all the newly synthesized DNA is found at the repair site, leaving the donor template unaltered ^2,3^. Additionally, both strands appear to be synthesized by DNA polymerase δ, which normally is confined to copying short, Okazaki-length segments on the lagging strand.

**Figure 1:**
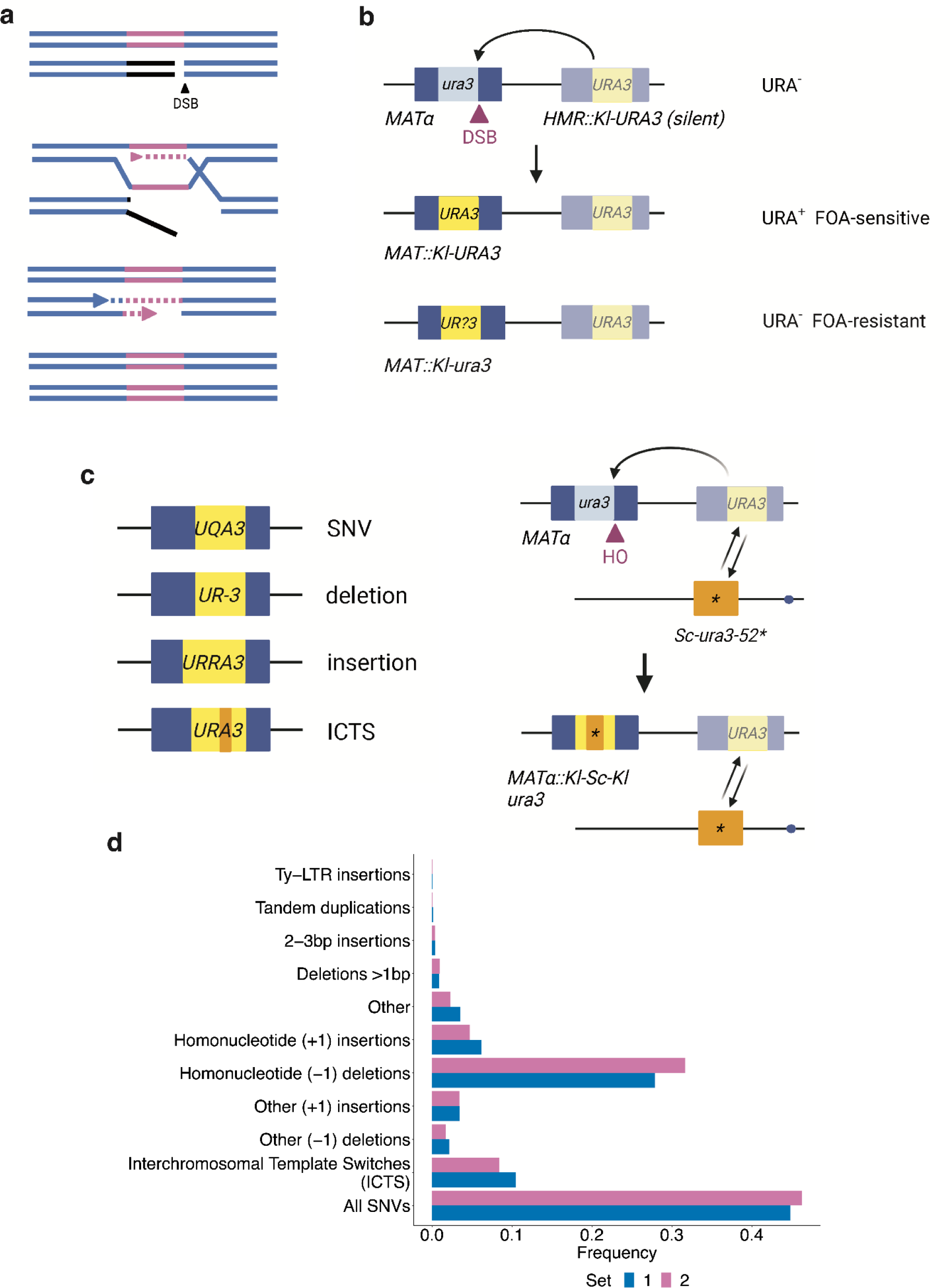
The spectrum of mutational events arising during repair replication. **a**, Schematic of synthesis dependant strand annealing (SDSA) used to repair DSBs induced at the *MAT* locus (pink) with the *HMR***a** doner region (blue). **b**, Repair of HO-induced DSB at the *MAT* locus repaired by copying from *HMR::Kl-URA3*. Growth on media lacking uracil and media containing 5-FOA is indicated. **c**, Types of mutational events observed, including ICTS events. **d**, Frequency of all mutational events observed during replication repair across two independent replicates

In our previous work, we studied differences between normal and repair replication by modifying the yeast mating type switching system. In this system, the inducible HO endonuclease creates a DSB at the *MAT* locus that is repaired via SDSA from nearby unexpressed homologous regions ^4^. By replacing the **a**1 coding sequences within one of these donor regions, *HMR***a**, with a *K. lactis URA3* gene (*Kl-URA3*), we could select for highly efficient switching events in which the *Kl-URA3* sequences are copied into the *MAT* locus, replacing Yα and expressing the *Kl-URA3* gene (**Figure 1b**) ^5^. After DSB induction and the 5’ to 3’ resection of the DSB ends, the right end of the DSB at *MAT* (*MAT-Z1*) strand-invades with the perfectly matched 230 bp of homologous sequences at *HMR*. Strand invasion is followed by the initiation of new DNA synthesis, with polymerase δ extending the invading strand by copying *Kl-URA3* sequences. In normal switching events, the newly copied DNA extends into the left homology region, *HML-X* and the displaced, newly copied strand pairs with the resected second end of the DSB (called second-end capture), allowing replacement of the original Yα sequences by *Kl-URA3*. However, if copying of *HMR::Kl-URA3* is interrupted, the partly copied sequence can apparently dissociate from its template and undergo a number of different mutagenic fates, in which the end of the dissociated strand can pair via short microhomologies to produce a variety of mutations that are recovered at *MAT* ^5^. Virtually all of these Ura^-^ mutants have alterations within the *Kl-URA3* sequences that were copied into the *MAT* locus, while the unexpressed donor sequence remains unaltered. That these mutations arose during switching can be demonstrated by un-silencing *HMR::Kl-URA3* in such mutants, by addition of the Sir2 inhibitor, nicotinamide, whereby Ura-cells become Ura^+^, as the *HMR::Kl-URA3* donor sequence is unaltered.

Compared to mutations arising spontaneously during normal DNA replication of the same sequences, the mutation rate accompanying DSB repair is 1000-fold higher and the mutants display a remarkably different spectrum. Among spontaneous mutations the great majority of events are base pair substitutions, whereas most repair-replication events appear to involve some sort of slippage or dissociation of the partly synthesized new strand. These include -1 frameshifts in homonucleotide runs, intragenic deletions, quasipalindrome mutations, and interchromosomal template switches (ICTSs) that we have documented previously ^5,6,7^. Most of these alterations involve the apparent use of microhomologies at the rearrangement junctions.

The most complex events – ICTS - requires two template “jumps” into and from a 72% identical *S. cerevisiae URA3* sequence (*Sc-ura3-52*) on a different chromosome. After a few hundred bases of *Sc-ura3* are copied, there must be a second template switch back to the original *Kl-URA3* template, where copying proceeds until the shared homology between *HMR* and *MAT* (the X region) is reached, so that these sequences can be captured at *MAT* and a second strand synthesis can complete the process. Despite their divergence at the DNA level, most jumps between *Kl-URA3* and *Sc-ura3-52* are in frame, at small microhomologies, so that most events produce a *Kl-Sc-Kl* chimera that is fully functional. Initially, we only recovered Ura^-^ ICTS mutants that misaligned the *Kl* and *Sc* sequences at one particular 5-bp out-of-frame microhomology, resulting in frameshifts ^5^. In subsequent work, we enriched for ICTS events by demanding correction of a 32-bp deletion in the *Kl-ura3* sequence via copying the deleted bases from *Sc-ura3-52* ^6^. The rate of ICTS is strongly limited by the degree of divergence. When the *Sc-ura3-52* sequences were replaced by *K. lactis* sequences (i.e. the jumps were between identical sequences, correcting the 32-bp deletion), the rate of ICTS increased more than 500-fold, to a level of > 1 in 10^3^ repair events ^6^.

Our prior work was based on the individual sequencing of independent events so we were only able to examine the sequence of approximately 50-100 mutational events per condition. Here we report PacBio sequencing of a large population of repair-associated mutations. From these larger samples we could identify many intragenic deletions and insertions, also bounded by microhomologies. We also assured recovery of most ICTS events by introducing a -1 frameshift into *Sc-ura3-52* so that most of these template switches would be Ura^-^.

This spectrum of mutations reflects the instability of the repair replication fork and its architecture. Specifically, we deduce that the D-loop at the donor containing the newly replicating strand is open ≥ 100 bases ahead of the DNA polymerase while it is rapidly closed behind the polymerase, as the donor strands reanneal. Further, our observation of many more -1 deletions than +1 insertions arising during repair suggests either differences in DNA polymerase δ activity during repair versus normal replication, or else reflect influences of the mismatch repair system on normal replication.

## Results

### Isolation of DSB repair-associated mutations

We isolated a large number of mutations arising after induction of *MAT*α switching using the transcriptionally silenced *HMR::Kl-URA3* donor as described above. Mutations were selected as Ura^-^ (5-FOA-resistant) colonies after galactose-induced recombination (**Figure 1b**). As noted above, ICTS events create chimeric Kl-Sc-Kl URA3 genes that are generally in-frame and functional. To obtain sufficient numbers of ICTS events in this context, we created a -1 frameshift mutation and a single base pair change in *Sc-ura3-52* so that any gene conversion that copied the middle segment of this donor template would create a Ura3-Kl-Sc-Kl chimera after switching to *MAT*, so that ICTS events that copied this region would then be readily recovered along with other 5-FOA-resistant outcomes such as base pair substitutions, -1 frameshifts, intragenic deletions and other events (**Figure 1c**). This donor is designated *Sc-ura3-52\**. PacBio sequencing of the PCR-amplified *MAT* locus of ∼50,000 independent FOA-resistant mutations that grew as 5-FOA-resistant colonies revealed the spectrum of mutations arising during repair replication (**Figure 1d**). We performed this experiment with two independent biological replicates and one technical replicate (the first biological replicate is noted as “set 1”, the technical replicates are averaged and notated as “set 2”. Numbers of events for each set are recorded in **Table S1**). While set 1 had more reads than set 2, the two largely showed the same spectrum of mutations.

Overall, these results agree with those obtained previously by sequencing individual colonies selected for mutational events ^5,6^. Approximately 50% of the Ura^-^ colonies arose from base-pair substitutions and 30% were -1 frameshifts, the great majority of which were in homonucleotide runs. A small fraction were ∼300 bp insertions that proved to be sigma sequences that are the LTRs of yeast retrotransposon Ty elements. About 10% of mutations were ICTS events we had observed in previous experiments. Finally, about 10% of mutations were +1 frameshifts, which were slightly more common in homonucleotide runs (64%). We did not search specifically for quasipalindrome mutations that are among the complex events listed as “other.”

### Repair replication leads to many more -1 than +1 frameshifts, in contrast to normal replication

As we had observed previously, a frequent class of mutations arising during DSB repair were -1 frameshifts, 93% of which were in homonucleotide runs. In contrast, +1 insertions were much less frequent, and 64% occurred in homonucleotide runs. Among mutations in homonucleotide runs, -1 were found 4.5 times more often than +1. To compare these results to the spectrum of +1 and -1 frameshifts among spontaneous mutations, we analyzed the whole-genome sequencing results obtained by Liu and Zhang (**Table S2**)^7^. Among the spontaneous mutations, 96% of all +1 and -1 mutations arose in homonucleotide runs. Moreover, during replication, the median number of +1 insertions in homonucleotide runs were almost equal to the median number of -1 deletions (**Figure 2a**). The strong bias towards -1 frameshifts in HO-induced DNA repair synthesis (**Figure 2b**) most likely reflects intrinsic differences in the DNA replication machinery present during normal chromosome duplication and that which is assembled to copy donor sequences during DSB repair.

**Figure 2:**
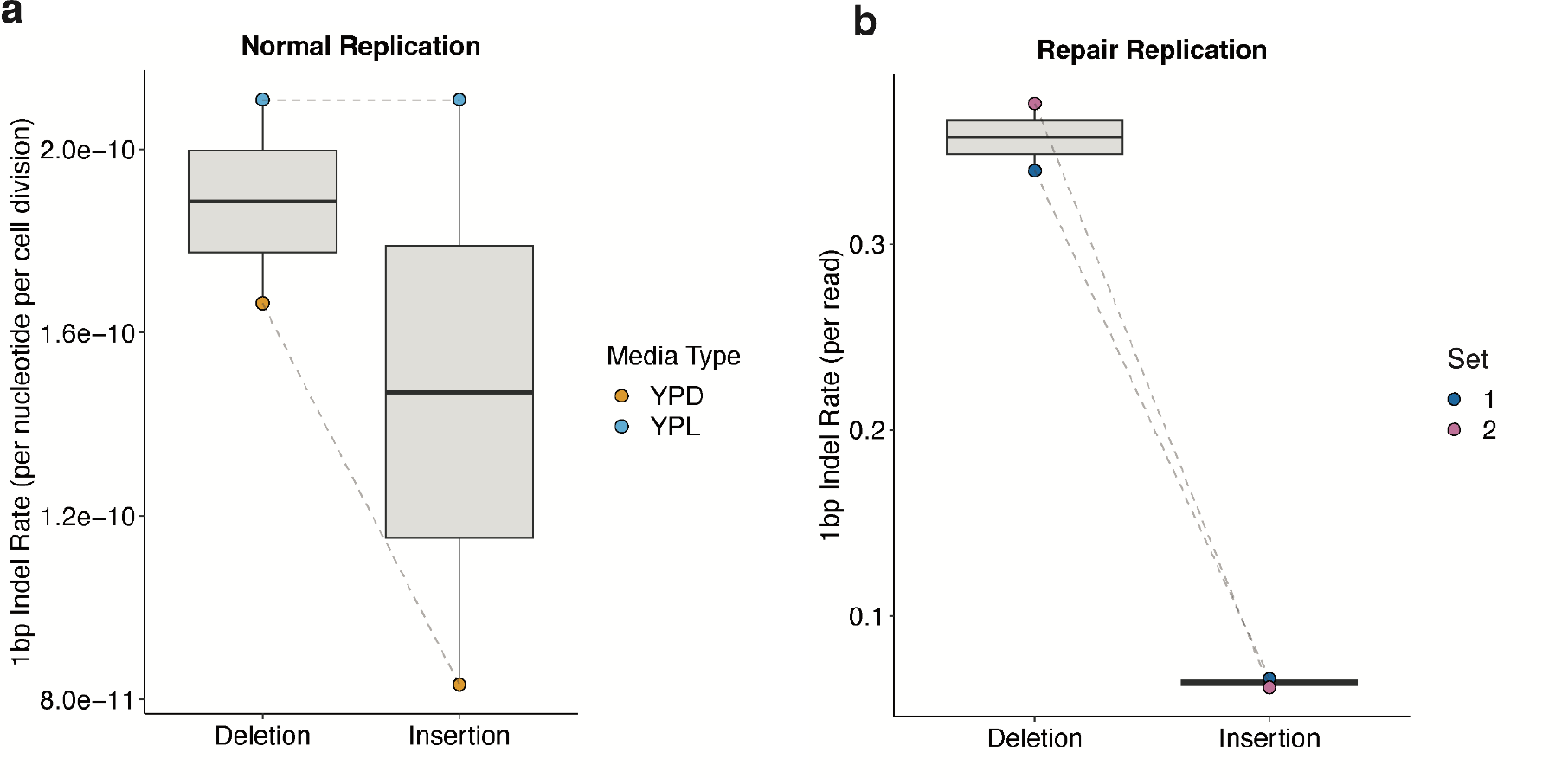
-1 indels are more common in repair replication than normal replication. **a**, Rate of 1bp insertions and deletions during normal replication on YPD and YPL media. **b**, Rate of 1bp insertions and deletions during repair replication.

### Interchromosomal template switches are frequent in replication repair, even with a highly divergent donor

As described above, we used the *Sc-ura3-52\** strain to enhance the recovery of interchromosomal template switches (ICTS). Even though the *Sc-ura3-52* sequence is only 72% identical to the *HMR::Kl-URA3* template that is initially engaged by the DSB end, 9% of all the Ura-events we recovered were ICTS events, >99.5% of which copied the -1 mutation into the chimeric sequence. The few ICTS events that did not include this region displayed some other frameshift alteration that accounted for the Ura-phenotype.

On average, ICTS events included 165 bp copied from *Sc-ura3-52\** (**Figure 3a**). Each of these “jumps” was bounded by a region of microhomology (MH) (on average 7.5 bp) shared between Kl-URA3 and *Sc-ura3-52\** (**Figure 3b**). A heatmap of the locations of the most frequently used MHs shows that the longest MHs on either side of the -1 mutation are most often used (**Figure 3c, S1**). However, some very short MHs (i.e. 2 bp) are used much more often than would be expected based on their length. Similarly, there are some long MHs (e.g. an 11-bp sequence underlined in **Figure 3c**) that are used 10 times less often than many 6-bp MHs. However, these under-utilized longer MH sequences are further from the location of the -1 mutation that allows recovery of the Kl-Sc*-Kl ura3 chimeras, so it is possible that their under-utilization reflects an intrinsic constraint of the length of DNA that can be copied during ICTS. We note that we define MH usage by the standard practice in the field, counting the number of matches until a single mismatch is encountered, so that contribution of additional MH further from the junction is discounted.

**Figure 3:**
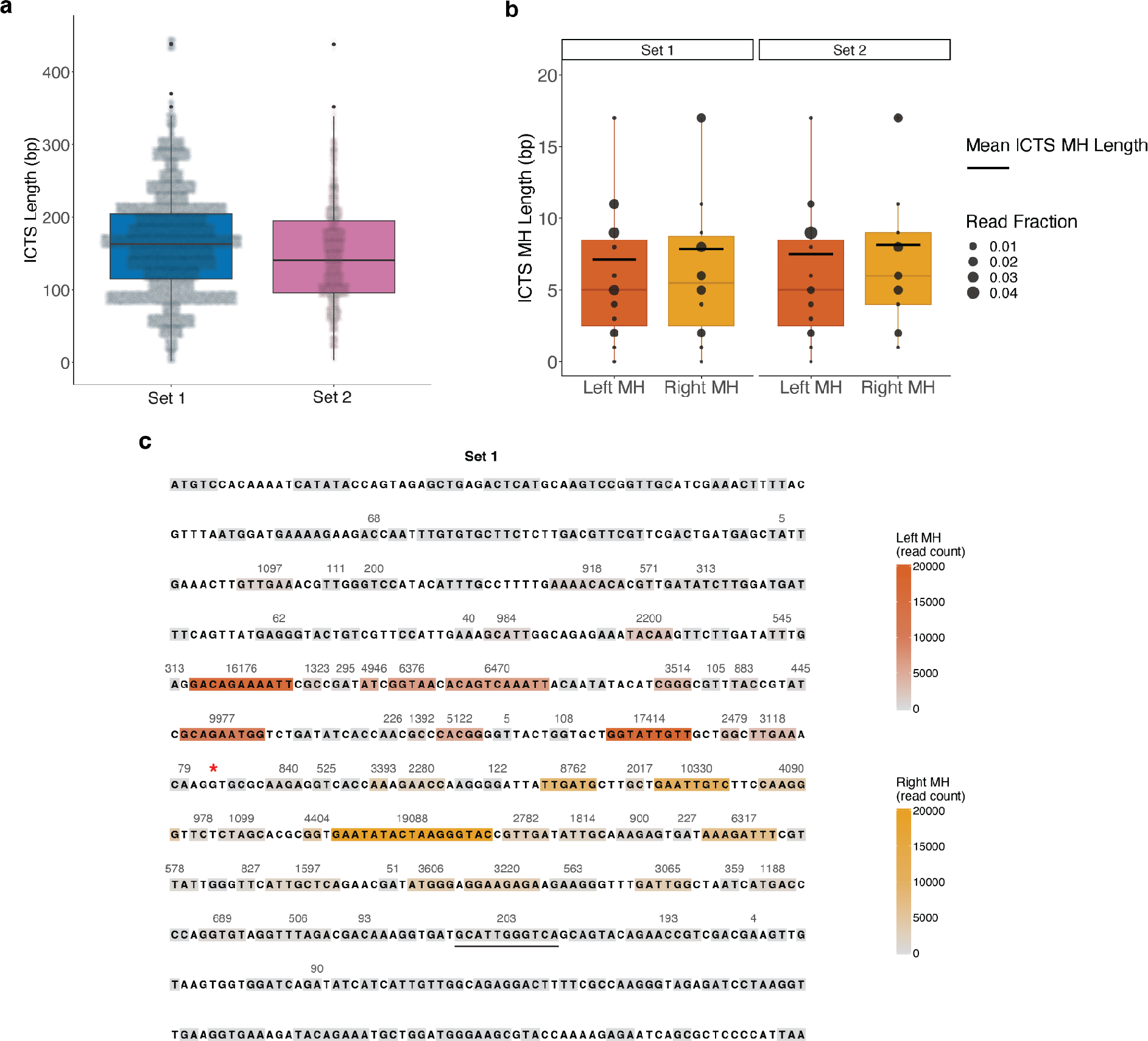
ICTS events are observed during repair replication. **a**, The length distribution of ICTS events observed during replication repair. **c**, The length distribution of MHs flanking ICTS events. **c**, The frequency with which each MH between *Kl-URA3* and *Sc-ura3-52\** was used for ICTS events. Grey boxes indicate unused MHs, orange boxes indicate MHs used to jump “in” to *Sc-ura3-52\**, and red boxes indicate MHs used to jump “out” of *Sc-ura3-52\**. Darker colors indicate more frequent usage. An 11bp MH used much less frequently than 6bp MHs is underlined. MH, microhomology.

### Microhomology-bounded intragenic deletions are more abundant than intragenic tandem duplications

Within the copied *MAT::Kl-URA3* sequences we identified both MH-bound deletions and tandem duplications. Deletions are approximately 5 times more abundant than tandem duplications (TDs) (**Figure 4a**) and distinctly different in size. Deletions were on average 60 bp long; however there were almost no deletions smaller than 25 bp. In contrast, TDs were quite short, averaging only 20 bp. Both deletions and TDs are usually bounded by MHs: deletions have a mean MH length of 6 bp while TDs have mean MH length of 3 bp (**Figure 4b**). These characteristics suggest a picture of the repair replication fork in which the polymerase can jump further forward than backward (**Figure 4c**, see Discussion).

**Figure 4:**
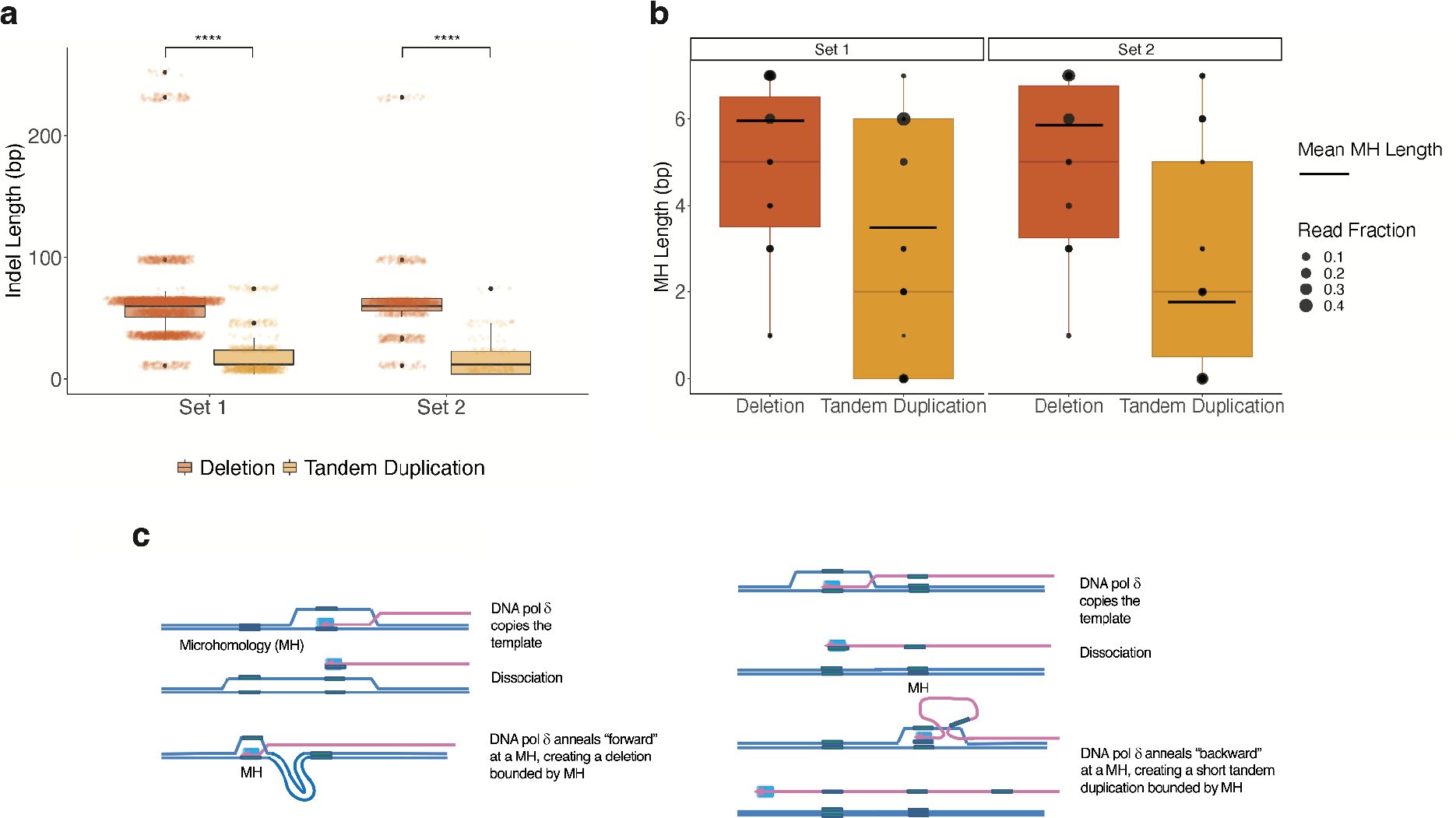
MH-bounded deletions are longer than tandem duplications. **a**, The length distribution of MH-bounded deletions and tandem duplications in repair replication. Deletions are significantly longer than tandem duplications, Wilcoxon Rank Sum test. P-values both < 2.2e-16 **b**, The length distribution of MH length for MH-bounded deletions and tandem duplications arising during repair replication. **c**, Schematic of the mechanism of formation of MH-bounded deletions (left) and tandem duplications (right). MH, microhomology.

### Microhomology-bounded intragenic deletions are not aided by additional adjacent microhomeology

We next set out to further characterize the nature of deletions arising during the copying of *HMR::Kl-URA3* into *MAT*. Using the *Sc-ura3-52* strain (without the -1 deletion), we isolated and pooled 50,000 FOA-resistant colonies and purified DNA. We then extensively digested the DNA with the *Bst*EII restriction endonuclease that cleaves one site in the *Kl-URA3* sequences and does not cleave in *ura3-52*. Thus only mutational events that remove the BstEII site, such as deletions and ICTS events, should remain. We sequenced the digested DNA by paired-end illumina sequencing. There were approximately 9 times as many deletions as ICTS events in this strain. Many different deletions were identified (**Figure 5b**). On average the deletions were 98 bp in length, bounded by MH of 6.2 bp. The frequencies of different deletions using the same number of adjacent MH varied widely; in the case of deletions with 5 bp MH, the most frequent events had >700,000 reads, while the least frequent were fewer than 100.

**Figure 5:**
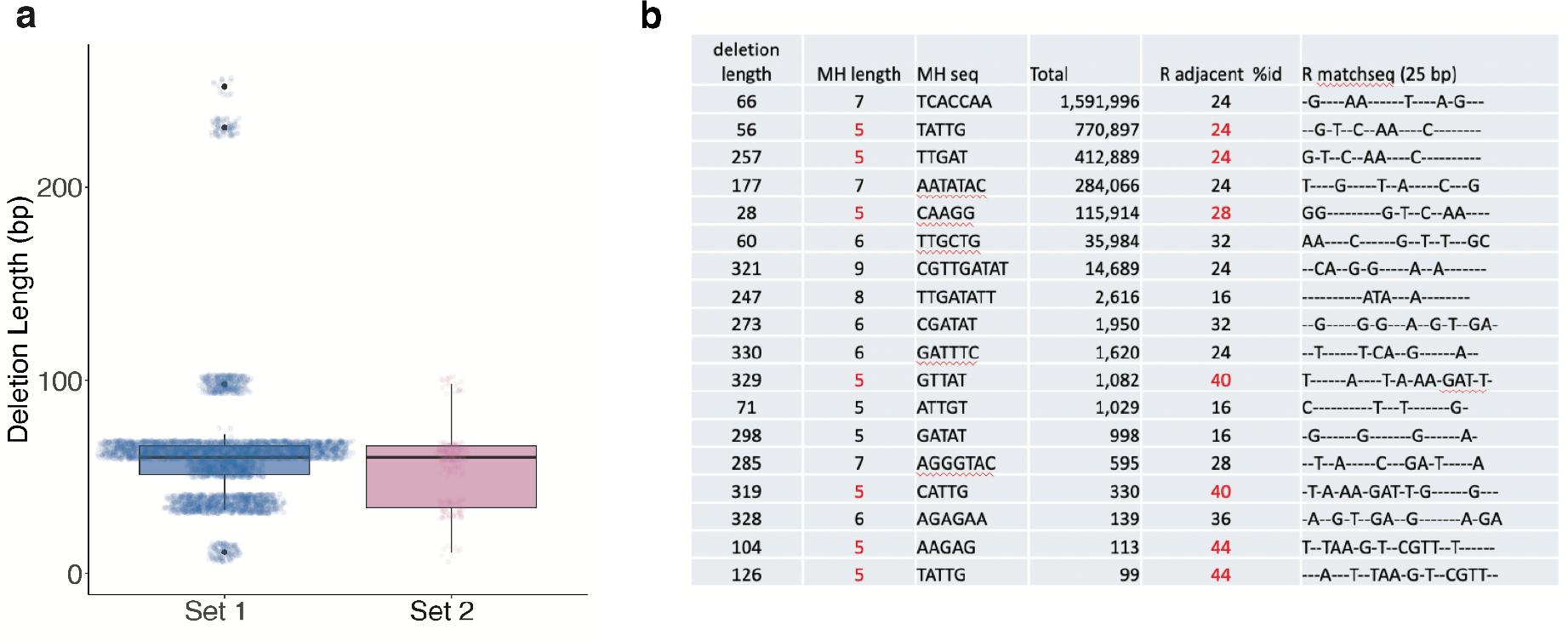
Homeology does not contribute to the choice of MH used for intragenic deletions. **a**, Length distribution of intragenic deletions observed without BstEII digestion. **b**, Comparison of homeology length, MH length, and frequency of MH usage across deletions identified with BstEII digestion. MH, microhomology

We entertained the hypothesis that the deletions used most frequently shared more adjacent homeology that could assist in stabilizing the intermediate leading to a deletion even though by conventional nomenclature these adjacent sequences were not part of the MH at the boundary. We calculated the percent base identity for the 25 bases upstream of the MH, to determine if more frequently used MHs had more homeology adjacent to the MH. However, as seen in **Figure 5b**, there was no strong correlation between the amount of homeology in the 25 bases upstream bases and their rate of usage; indeed several of the least used sequences had the highest fraction of adjacent base pair matches in the 20 bp upstream of the MH.

## Discussion

By sequencing a large number of structural variants arising as mutations during DSB repair, we have identified several aspects of the repair process that are distinctive for repair-induced mutations compared to those arising during simple replication of the same sequences.

As we have shown previously, a large proportion of the mutation repair events involve some form of dissociation of the partly-copied strand that will become assimilated at the recipient locus. These dissociations enable the formation of intragenic deletions and tandem duplications, quasipalindromes, and interchromosomal template switches. The simplest example of this “slippage” are -1 frameshifts in homonucleotide runs, which greatly outnumber +1 insertions. In normal replication -1 and +1 frameshifts occur at equal rates. This difference makes clear that the repair replication apparatus differs in distinct ways from the normal replication fork. In normal replication, leading and lagging-strand synthesis are coordinated and apparently closely associated with proofreading by the MSH-MLH mismatch repair system. However, DSB repair using synthesis-dependent strand annealing occurs in two separate steps. A first strand is copied and only later – after second end capture – is the first strand used as a template to complete the repair event^9^. Moreover, the mismatch repair machinery appears to play little or no role in correcting the errors arising during DSB repair, possibly because this proofreading apparatus does not follow closely behind the repair replication fork as it apparently does when there is a complete simultaneous leading- and lagging-strand replication fork^5^. The many more -1 deletions arising during repair may reflect some intrinsic feature of DNA polymerase δ when it is not part of a coordinated replication fork; alternatively the ratio of +1 and -1 events may be influenced during normal replication by the mismatch repair system. That DNA polymerase δ is the principal DNA polymerase used in repair is evident from the fact that a proofreading-defective mutation (*pol3-01*) of Pol δ dramatically reduces the appearance of all classes of structural variants in our repair system^5^. We attribute the absence of the frequent slips and jumps to the observation that a proofreading-defective Pol δ is more “processive” and less likely to dissociate from its template, as demonstrated *in vitro*^10^.

The pattern of MH-bound deletions and tandem duplications that we observe suggests a picture of the repair replication fork (**Figure 6**). After the DSB end is resected and bound by Rad51, strand invasion occurs at the region of homology on the right side of *HMR::Kl-URA3*, creating a D-loop. (In the *MAT* switching system, the presence of 700-bp nonhomology to the left of the HO cleavage site biases the initial strand invasion to use the *HMR-Z* region^8^. The 3’ end of the invading strand then is used by DNA Pol δ to copy the donor, by a moving replication “bubble” in which the region ahead of the fork opened either by the movement of the polymerase itself or by a helicase; however we have previously shown that gene conversion events such as *MAT* switching are not dependent on the normal replicative CMG helicase^11^. We suggest that the D-loop is kept open by the binding of the single-strand DNA binding protein complex, RPA. An intrinsic feature of this repair process is that the newly-copied strand does not remain semi-conservatively bound to its template; rather the strand is displaced so that eventually all the newly-copied DNA is found at the recipient locus. Inherent in this process, the previously copied region behind the fork soon “closes up.” Because of the displacement of the new strand, the repair replication fork is unstable; indeed if a helicase driving the new-strand displacement is more active than the rate of polymerization, the newly-copied strand may be completely dissociated. Such dissociations are necessary for events such as ICTS, quasipalindrome mutagenesis and both deletions and tandem duplications. Dissociation may also be responsible even for the slippage in -1 frameshifts.

**Figure 6:**
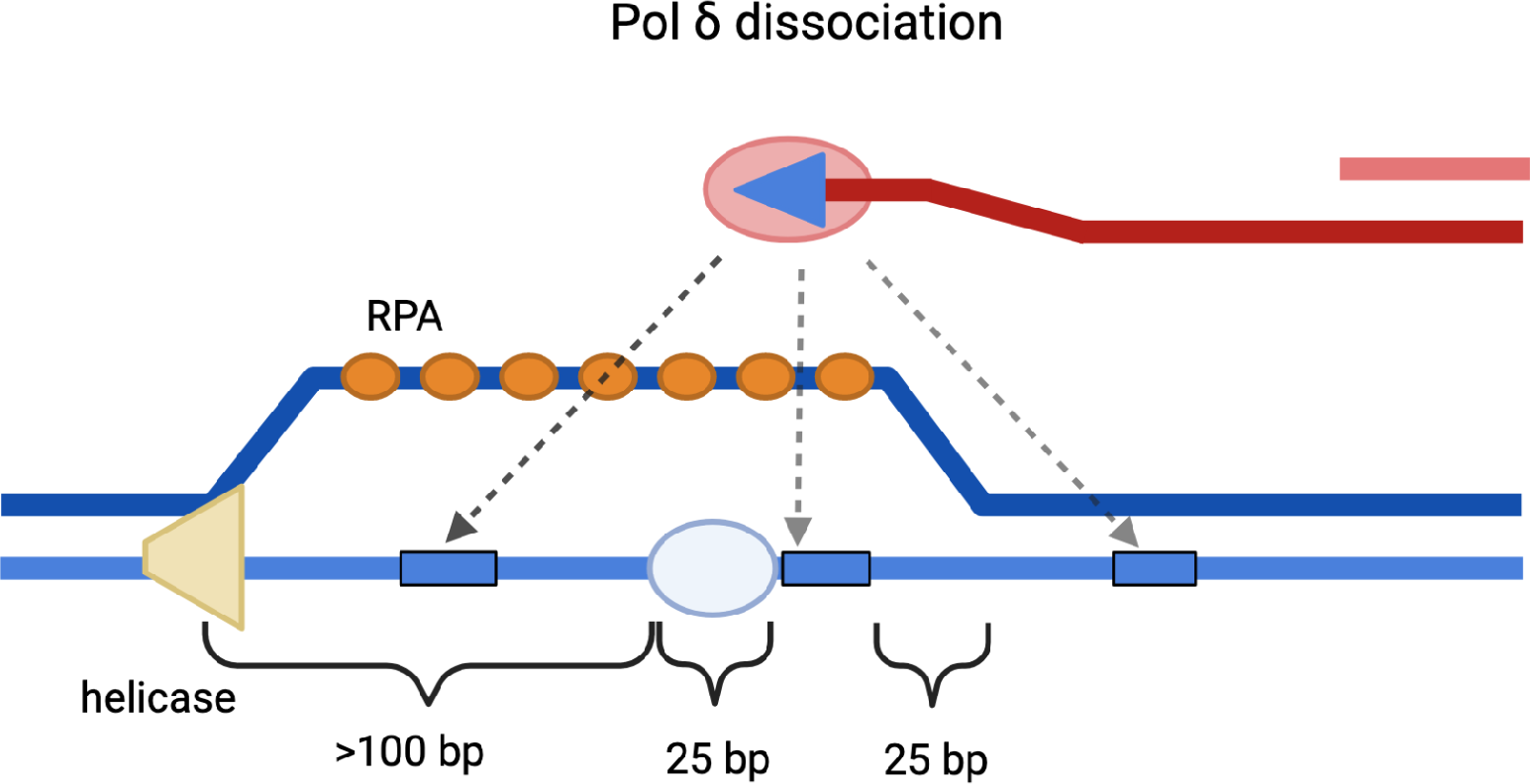
Schematic of the repair replication fork. A helicase precedes the migrating D-loop >100 bp ahead of the polymerase. An unidentified component of the repair replication machinery occupies about 25 bp immediately ahead of the polymerase.

We observed that deletions are longer than tandem duplications, and that there are few deletions shorter than 25 bp. Based on this pattern we suggest that the dissociation of the partly-copied strand, likely still attached to DNA polymerase δ, leaves behind PCNA and perhaps other factors still bound to the template strand, preventing annealing in that region. Further we suggest that the creation of deletions or tandem duplications requires that the template strand remain single-stranded and accessible to polymerase-mediated annealing either in front of the point of dissociation (creating a deletion) or behind that point (creating a tandem duplication) (**Figure 4c**). The presence of RPA in the non-template strand should help maintain this open structure, but the sequences behind the fork will likely be “closed up” as the strand is dissociated, leaving only a very short region where a tandem duplication might be generated. This picture of the pair fork is supported by *in vitro* and *in vivo* mapping of the size of a D-loop generated by Rad51 in yeast^12^.

Further refinement of this picture may be gained by examining the consequences of mutations that alter the processivity of DNA polymerase or various helicases as well as factors associated with loading and unloading PCNA and other replication factors. This investigation is ongoing.

Many aspects of this dissociation are not known. Does the strand become detached from the DNA polymerase? Does the DNA polymerase become detached from PCNA and does PCNA remain bound to the template? Do ICTS or other MH-bounded events require the participation of Rad51? Because Rad51 is required for the initial strand invasion, it is difficult to assess its role in secondary invasion events. Further experimentation with our *HMR::Kl-URA3* system will yield important insight into the mechanics of and participants in repair replication.

## Materials and Methods

### Yeast Strains and Plasmids

All strains were derivatives of strain WH50 ^5^ with genotype: *ho hml::ADE1 MAT*α *hmr::Kl-URA3 ade1 leu2 lys5 trp1::hisG ura3-52 ade3::Gal-HO*.

Strain QW904 was modified by Cas9-mediated gene editing of *ura3-52* to delete G409 and change G411 to C so that the otherwise in-frame open reading frame contained a marked 1-bp deletion.

### Isolation of Mutations During DSB Repair

Approximately 10^7^ cells were plated on YEP-GAL to induce the expression of HO endonuclease. After overnight growth, these plates were replica-plated to FOA. Because the mutation rate associated with DSB repair is approximately 1000-fold higher than spontaneous mutations, each FOA-resistant colony that arises likely represents an independent repair-replication associated event. FOA-resistant colonies were purified by further replica-plating on FOA plates and then either individual colonies were isolated for individual sequencing or several hundred FOA-resistant colonies per plate were pooled among 100 plates (approximately 11,000 independent colonies) and DNA was isolated from the pool. A larger set of FOA-resistant mutants were isolated from strain WH50.

DNA specific for the region including *MAT* was purified by PCR using primers specific for *MAT::Kl-URA3* and not the *HMR::Kl-URA3* donor (**Table S3**). Pooled samples were sequenced by PacBio across a 1.6-kb region.

### Alignment and mutation calling of *MAT::Kl-URA3* sequence from Ura^+^ Colonies

Due to the presence of ICTS events, alignment to *sc-URA3* was impossible with existing methods, preventing calling of mutations. Therefore, we created a custom alignment algorithm based on Smith-Waterman^13^ and Breakpointer methods^14^.

First, to determine if each sequence contains an ICTS event, they were aligned to *Kl-URA3* using a global Needleman-Wunsch algorithm using the following scores:

| Initial Alignment |  |
| --- | --- |
| Match | 1 |
| Mismatch | -80 |
| Gap open | -500 |
| Gap extend | -10 |

The score of this alignment was normalized to the length of the aligned query (as a proxy for highest possible score). Alignments with a normalized score higher than -2 were kept. Alignments with a lower normalized score were passed on to the modified alignment algorithm that allows jumping between *Kl-URA3* and *Sc-ura3-52*. All scores and thresholds were determined empirically. In all alignment matrices, the sequence to be aligned is along the top of the matrix (aka in the columns) and the reference sequence (*Kl-URA3* or *Sc-ura3-52\**) is along the side (aka in the rows).

The first step of the modified alignment algorithm is to do a global/local alignment of each sequence to *Kl-URA3*. To force inclusion of the left side of Kl-URA3, the alignment matrix is initialized as a Needleman-Wunsch matrix, with 0 only at [1,1], and gap penalties applied for all other [1,] and [,1] cells. Scores for this alignment, called M1, are set to optimally find the best matching left-most chunk of each sequence, so it has moderate mismatch score, but high gap scores:

| M1 Alignment |  |
| --- | --- |
| Match | 2 |
| Mismatch | -30 |
| Gap open | -40 |
| Gap extend | -90 |

After finding the best left-most alignment between each ICTS-containing sequence and *Kl-URA3*, we do a local Smith-Waterman alignment between each sequence and *Sc-ura3-52\**. Since this will be the middle of the final alignment it is not required to include all of the sequence, so the alignment matrix is initialized with 0’s in [,1] and [1,]. Now mismatch, deletion and insertion scores are lowered to allow for some errors within the gene conversion. Additionally, this alignment, called M2, is also allowed to ‘jump’ back to the M1 alignment matrix (ie, template switch to *Kl-URA3*). Specifically, at each cell of the alignment matrix, there is an option for the alignment to jump to the maximum cell in the previous column of M1. This corresponds to keeping the same index of the sequence, but allowing a jump to anywhere in *Kl-URA3*. M2 does not allow for affine gaps.

| M2 Alignment |  |
| --- | --- |
| Match | 2 |
| Mismatch | -6 |
| Deletion | -20 |
| Insertion | -3 |
| Jump to M1 | -1 |

This step recovered sequences that were erroneously rejected due to the strictness of the cutoff for accepting the Inital alignment. These sequences lack ICTS events, so the M2 alignment will not extend very far beyond the M1 alignment. Therefore, any M2 alignments that extend M1 less than 15 bases are rejected in favor of the Initial alignment.

If the M2 alignment is accepted, the last step is to create an M3 alignment of the sequence to Kl-URA3, allowing jumping back to M2. To keep the location of the jump between M1 and M2, we first create a matrix to combine M1 and M2 into one alignment matrix to allow M3 to jump back to the previously completed Kl-Sc alignment. Because we are constructing a reference for this intermediate “new reference” alignment, this alignment is between the sequence and the aligned reference sequence from M2. This new alignment matrix needs to have a higher match score to encourage the third and final alignment to jump to it. That correspondingly requires increasing the other scores as well:

| MNewRef Alignment |  |
| --- | --- |
| Match | $805 = \text{nchar}(\textit{KI-URA3}) + 1$ |
| Mismatch | $-67 = -1 * \text{nchar}(\textit{KI-URA3}) / 12$ |
| Deletion | $-844 = -1 * (\text{nchar}(\textit{KI-URA3}) + 40)$ |
| Insertion | $-844 = -1 * (\text{nchar}(\textit{KI-URA3}) + 40)$ |
| Jump to M2 | -1 |

This alignment is initialized to force inclusion of the left-most edge of the query and reference (initialized as a Needleman-Wunsch matrix, with 0 only at [1,1], and gap penalties applied for all other [1,] and [,1] cells). Traceback starts at the max cell in the bottom row, forcing inclusion of the entire M2 reference alignment.

The final alignment is then constructed with the M3 alignment between the sequence and *Kl-URA3*, allowing jumping to the MNewRef reference alignment (which is essentially the *Kl-URA3* and *Sc-ura3-52\** reference sequences pasted together at the first microhomology of each sequence’s ICTS event). This is a global alignment, specifically the M3 alignment matrix is initialized initialized as a Needleman-Wunsch matrix, with 0 only at [1,1], and gap penalties applied for all other [1,] and [,1] cells and the traceback is initialized at the bottom right cell, to force inclusion of the entire reference and query sequence. Scores for this matrix are lower, to allow jumping back to MNewRef & tolerance of any SNVs:

| M3 Alignment |  |
| --- | --- |
| Match | 1 |
| Mismatch | -3 |
| Deletion | -10 |
| Insertion | -10 |
| Jump to MNewRef | -1 |

Mutations were identified by comparing between the aligned sequences resulting from this final alignment or the Initial alignment and the reference *Kl-URA3* and *Sc-ura3-52\** sequences.

### Statistical Methods

Analyses were performed in R version 4.2.1.

### Enrichment of Intragenic Deletions

We generated 50,000 independent FOA-resistant, *MAT::Kl-ura3* mutations in the *Sc-ura3-52* strain, pooling the colonies and subjecting the purified DNA to digestion by BstEII-HF, which cleaves a site in the middle of the *Kl-URA3* sequence. In this way, most SNVs, -1 frameshifts, TDs and many other events that retained the BstEII site were eliminated, thus enriching for deletions that removed this restriction endonuclease site. ICTS events that replaced the region around BstEII-HF with the divergent sequences from *Sc-ura3-52* were also recovered. The resulting DNA was PCR amplified using a *MAT*-specific and a *Kl-URA3*-specific primer and then subjected to paired-end Illumina sequencing. These sequences were analyzed as described above.

## Acknowledgments

Initial DNA sequencing and analysis was performed by Dr. Alex Ferrazzoli, who passed away in 2020. Figures 1A-C and 6 were created with BioRender.com. We thank and acknowledge the following funding sources: the Gray Matters Brain Cancer Foundation (R.B.), the Pediatric Brain Tumor Foundation (R.B.), and Break Through Cancer (R.B.). Research in the Haber lab is supported by NIH grant R35 GM127129. Summer research support for V.L. was through a Frederick W. Alt Biology Summer Research Fellowship. S.D. is supported by an NIH NRSA award (F32), 1F32CA261024.

## Author contributions

N.S., Q.W., T.C., and V.L. performed the experiments. S.D., S.W., and S.Z. analyzed the data. J.E.H. and S.D. wrote the manuscript with input from all authors. S.D., R.B., and J.E.H. conceived the project and designed the experiments and analyses.

## Figures & Figure Legends

**Figure S1:**
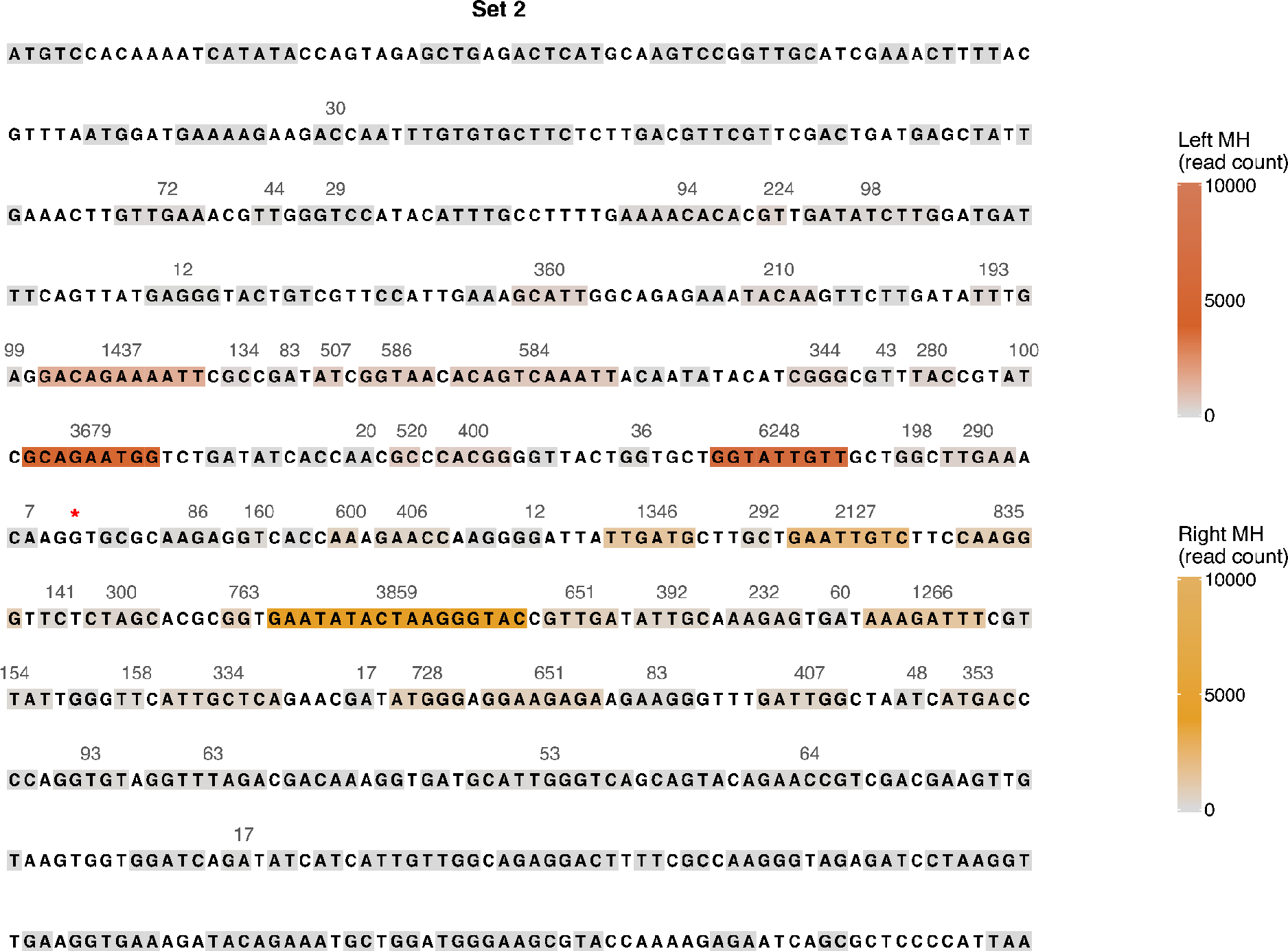
Frequency of usage of MHs between *Kl-URA3* and *Sc-ura3-52\** to form ICTS events in set 2. Grey boxes indicate unused MHs, orange boxes indicate MHs used to jump “in” to *Sc-ura3-52\** and red boxes indicate MHs used to jump “out” of *Sc-ura3-52\**. Darker colors indicate more frequent usage. MH, microhomology.

